# Cross-species systems analysis of evolutionary toolkits of neurogenomic response to social challenge

**DOI:** 10.1101/219444

**Authors:** Michael C. Saul, Charles Blatti, Wei Yang, Syed Abbas Bukhari, Hagai Y. Shpigler, Joseph M. Troy, Christopher H. Seward, Laura Sloofman, Sriram Chandrasekaran, Alison M. Bell, Lisa Stubbs, Gene E. Robinson, Sihai Dave Zhao, Saurabh Sinha

**Author notes:** To whom correspondence should be addressed. Address correspondence to: Sihai Dave Zhao:; Saurabh Sinha.

## Abstract

Social challenges like territorial intrusions evoke behavioral responses in widely diverging species. Recent work has revealed that evolutionary “toolkits” – genes and modules with lineage-specific variations but deep conservation of function – participate in the behavioral response to social challenge. Here, we develop a multi-species computational-experimental approach to characterize such a toolkit at a systems level. Brain transcriptomic responses to social challenge was probed via RNA-seq profiling in three diverged species – honey bees, mice, and three-spined stickleback fish – following a common methodology, allowing fair comparisons across species. Data were collected from multiple brain regions and multiple time points after social challenge exposure, achieving anatomical and temporal resolution substantially greater than previous work. We developed statistically rigorous analyses equipped to find homologous functional groups among these species at the levels of individual genes, functional and coexpressed gene modules, and transcription factor sub-networks. We identified six orthogroups involved in response to social challenge, including groups represented by mouse genes *Npas4* and *Nr4a1*, as well as common modulation of systems such as transcriptional regulators, ion channels, G-protein coupled receptors, and synaptic proteins. We also identified conserved coexpression modules enriched for mitochondrial fatty acid metabolism and heat shock that constitute the shared neurogenomic response. Our analysis suggests a toolkit wherein nuclear receptors, interacting with chaperones, induce transcriptional changes in mitochondrial activity, neural cytoarchitecture, and synaptic transmission after social challenge. It reveals systems-level mechanisms that have been repeatedly co-opted during evolution of analogous behaviors, thus advancing the genetic toolkit concept beyond individual genes.

## INTRODUCTION

A pivotal idea arising from evolutionary developmental biology is that across the bilaterian clade, the same signaling and transcription factor genes, known as “toolkit” genes, underlie the patterning of basic morphological features such as the body plan and the eye (Carroll *et al*. 2005). This provides a conceptual framework for increasingly detailed explanations of developmental patterning in specific model organisms (Wilkins 2002). Moreover, its success has motivated researchers to ask if the toolkit framework, where common genetic programs coordinate fundamental processes and undergird shared phenotypes, is also applicable to studies of behavior (Rittschof & Robinson 2016; Toth & Robinson 2007).

Studying toolkits for behavior poses numerous challenges including: the relative paucity of detailed and directly comparable genetics and genomics datasets for behaviorally relevant phenotypes in most animal species, difficulties in defining correspondence between behaviorally relevant phenotypes in diverged species from different ecological contexts, and ambiguity regarding brain regions and other tissues where shared behaviorally relevant molecular mechanisms may manifest. Further, behaviors, being transitory and directly observable only while an animal is living, cannot be as readily gleaned from fossils as developmental phenotypes (Chen *et al*. 2013), giving us little evidence from the distant past that contextualizes what we observe in extant species.

In an example of an evolutionary approach, our group recently studied whether shared gene expression correlates constitute a toolkit for the neural response to a territorial intrusion by a conspecific – more generally referred to as a social challenge – in the mouse, the three-spined stickleback fish, and the honey bee (Rittschof *et al*. 2014). These three highly diverged model social species have well-assembled genomes, providing ample technical resources for detailed comparisons of functional genomic correlates. Further, phylogenetic analyses strongly suggest convergent evolution of relatively sophisticated social phenotypes for these species (Kapheim *et al*. 2015; Woodard *et al*. 2011). Though a toolkit would be derived within each individual species, it would contain a common core of conserved components important for coordinating brain response to social contexts, and an argument for such a toolkit requires the identification of shared functional correlates. Accordingly, we found robust transcriptional responses in brain gene expression profiles across these three species 20-30 minutes after exposure to the intruder and discovered several common molecular mechanisms associated with the intruder response. Though similar to evolutionary development studies in its pursuit of a “toolkit”, our earlier study was notably different for its use of gene expression as the primary means to identify toolkit genes rather than direct or indirect measures of gene sequence. Similarly, other groups have discussed conservation in the transcriptional correlates of aggressive behavior within the vertebrate subphylum (Freudenberg *et al*. 2016; Malki *et al*. 2016) and in arthropods (Asahina *et al*. 2014).

The success of the above studies in identifying shared mechanisms motivates a concerted effort towards more comprehensive and rigorous descriptions of behaviorally relevant evolutionary toolkits. However, further progress has been limited due to two factors. First, the prior studies measured expression at only a single time point after animals were exposed to the social challenge and relatively soon after exposure. Such a design cannot capture longer-acting genetic programs. This simple design also limits the power of this previous study to detect responses whose anatomical and temporal profiles are shaped by the unique cell biology, neuroendocrine, and metabolic properties of brains in these three species (Bukhari *et al*. 2017; Saul *et al*. 2017; Shpigler *et al*. 2017b).

Second, evolutionarily shared mechanisms are likely to be found at various levels of organization that are not reducible to single genes, which have been the main level of comparative analyses thus far. Shared mechanisms embodied by gene orthogroups comprising multiple orthologs and paralogs, coexpressed modules, groups of genes dedicated to specific known biological processes, or regulatory sub-networks (Rittschof & Robinson 2016) have eluded discovery so far. Analytic tools that can identify such higher order functional entities across multiple species, brain regions, and time points in the face of complex gene orthology relationships among highly diverged species have been lacking.

We report here the results of a detailed investigation of the shared molecular roots of social behavior, specifically neural response to social challenge, that remedies the above issues. We use both a powerful experimental design and a new suite of computational tools to identify mechanisms that are deeply shared across species at different levels of organization. For simplicity we will refer to these as homologous functional groups (HFGs); see Figure 1.

The experimental design integrated datasets on mice, sticklebacks, and honey bees. Insights into the social neural transcriptomes for each of these individual species have been published (Bukhari *et al*. 2017; Saul *et al*. 2017; Shpigler *et al*. 2017b) and the transcriptomic data deposited in public databases (see **Materials and Methods**). However, these datasets were sequenced in parallel, allowing for comparative analysis and discovery of shared mechanisms with minimal technical effects. Each dataset compared the neural transcriptomes of animals exposed to the social challenge of a conspecific resident-intruder with control animals exposed to a novel non-social stimulus of approximately similar size. Recording transcriptional events in a time series (30 min, 60 min, and 120 min after the exposure to the social challenge within each species) generated a holistic view of a dynamic transcriptional process while allowing for inter-species differences in the timing of transcriptional trajectory. These experiments probed discrete socially relevant brain regions in each species, which increased the specificity of RNA-seq signals relative to sequencing whole brains as was done previously (Rittschof *et al*. 2014) while making no assumptions about whether these brain regions are homologous across species. Altogether, this experimental approach permits the elucidation of common molecular correlates of social behavior that may not seem important in individual species but rise to significance when looking at all three species together.

The main goal of the present work is to integrate these cross-species data for overall comparative analysis and discovery of shared mechanisms. Such an approach allows the elucidation and unification of common molecular correlates of social behavior that may not seem important in individual species but rise to significance when looking at all three species altogether. We developed computational methods that allowed us to ascertain not only individual genes, but also coordinately expressed ontology terms, coexpression networks, and transcriptional regulatory cascades commonly associated with a behaviorally relevant stimulus across these distantly related species. Our work goes beyond existing cross-species studies of tissue-specific (Lin *et al*. 2014) or developmental time-course transcriptomes (Gerstein *et al*. 2014); here, we not only identify shared transcriptomic patterns but also rigorously test and quantify their associations with brain responses while accounting for complex homology relationships between and within species. We also address open computational problems uniquely associated with a comparative systems biology study such as ours: identifying shared coexpression modules and transcriptional regulators in diverged species, as well as performing functional annotations of gene modules in a cross-species manner that accounts for available gene orthology information.

We report below the discovery of significant shared mechanisms at varying levels of molecular organization, later discussing our conclusions from the aggregate of evidence at multiple levels of abstraction from genes to systems.

## MATERIALS AND METHODS

### Social Challenge Exposure, Sequencing, and Identification of Differentially Expressed Genes

The results described derive from three separate experiments that proceeded in parallel previously in three different species. Briefly, within each species, animals were exposed to either a conspecific resident-intruder (challenged) or a neutral non-social stimulus of roughly equal size and shape (control). For details about the specific experimental paradigms used within each individual species, see our previous work (Bukhari *et al*. 2017; Saul *et al*. 2017; Shpigler *et al*. 2017b). After exposure to either challenge or control stimulus, we waited either 30 min, 60 min, or 120 min to collect brains for transcriptomic data within each species.

RNA-seq data were collected as described previously (Bukhari *et al*. 2017; Saul *et al*. 2017; Shpigler *et al*. 2017b). The data for these three sets are deposited in the GEO under accession numbers: GSE85876 (honey bee), GSE80346 (mouse), and GSE96673 (three-spined stickleback). In each species, we used a 1 CPM cutoff for an equivalent of the smallest group size for expression, as proposed in the edgeR documentation (Robinson *et al*. 2010). DEGs for each time point compare transcriptomes of challenged (experimental) to neutral (control). FDR thresholds from each individual species paper – 5% for bee, 10% for mouse, and 10% for stickleback – were used to compile the DEG lists (**Figure 1A**). We chose a lower FDR threshold for the honey bee because its experimental design was more powerful and therefore produced more DEGs.

**Figure 1:**
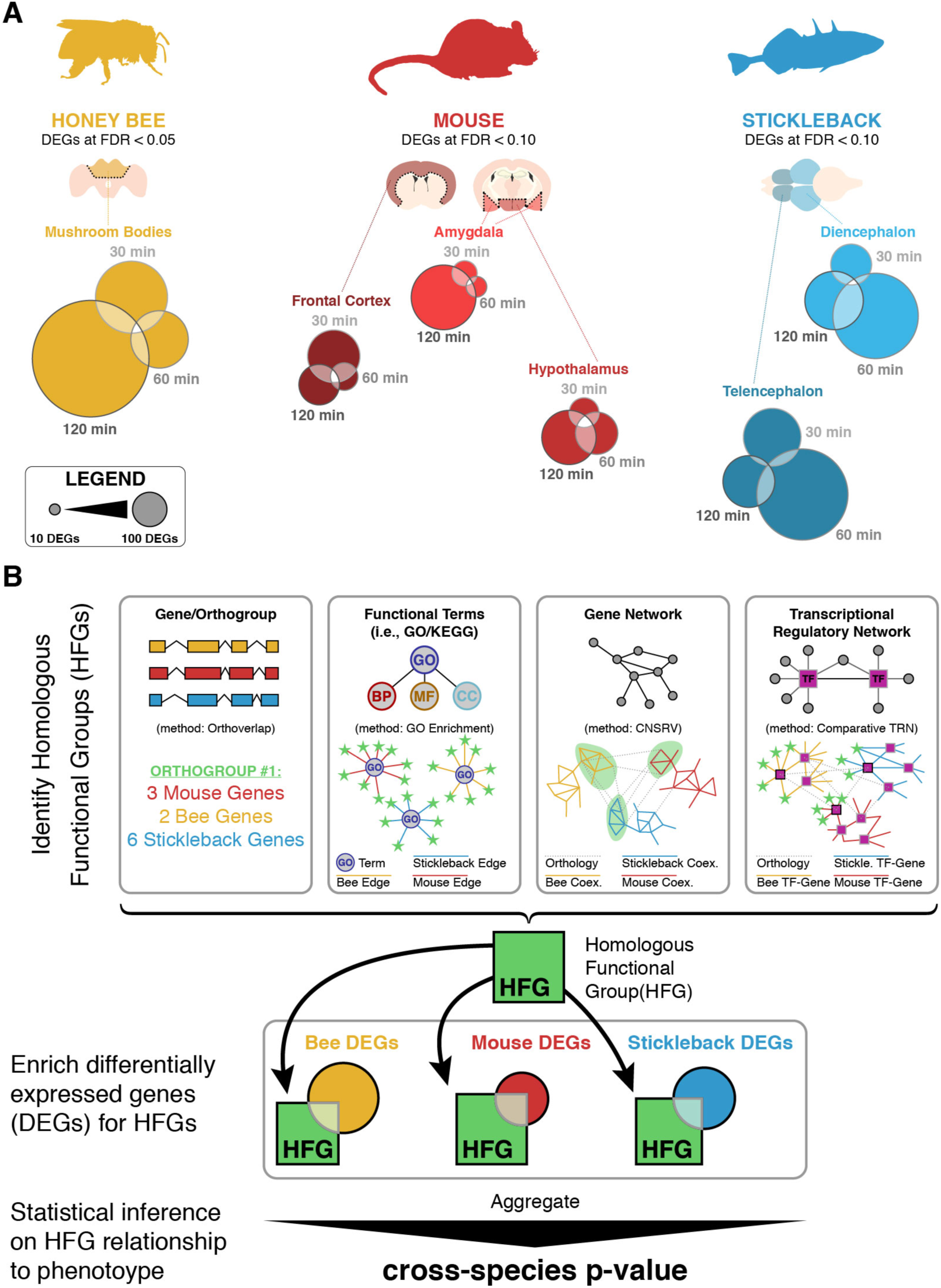
Summary of (**A**) dataset and (**B**) methods used within the paper to assess homologous function groups (HFGs). **A**) Description of dataset constituting multiple calls of differentially expressed genes (DEGs) within each brain region assayed for each species. Honey bee DEGs are called at FDR < 0.05 while mouse and stickleback DEGs are called at FDR < 0.10. **B**) Schematic of HFG analysis. Multiple methods of HFG identification utilized included: genes and orthogroups, functional terms, gene networks, and transcriptional regulatory networks (TRNs) and enrichment. HFGs are highlighted in green. Once HFGs were identified, enrichment of DEGs swithin each species was assessed prior to calculation of cross-species simultaneous enrichment.

### OrthoDB

Comparing DEGs between species requires a reliable orthology map between these three species. Using the raw data from OrthoDB v8 (Kriventseva *et al*. 2015), we first filtered for the three species of interest. We then identified the metazoan level orthogroups present within all of the three individual species used in this experiment, a total of 4,982 orthogroups. We found all paralogs inside of each orthogroup for the individual species, which brought us to a total of 10,158 genes in mouse, 6,725 genes in bee, and 10,869 genes in stickleback. The scripts used to annotate these orthogroups have been uploaded to GitHub (https://github.com/msaul/three_species_orthology).

### Multi-scale characterization of conserved molecular basis for analogous cross-species phenotype

We probed for an evolutionary toolkit for social challenge response at multiple levels of molecular organization in a uniform and systematic manner. For each level of organization – individual genes, cellular processes, coexpression modules, and TF regulons – we first identified HFGs (**Figure 1B**) in the three species as sets of genes that exhibited intra-species as well as inter-species commonality, e.g., involvement in the same cellular process, being paralogs or orthologs of each other, etc. We then tested each HFG for association with phenotype across all three species (see below). This systematic two-step approach is a novel feature of our work, and while our previous work (Rittschof *et al*. 2014) reported an initial use of the approach, it is developed fully in this work with a focus on statistical rigor.

### Identifying orthogroups with a conserved response to social challenge

We identified a given HFG as associated with phenotype if its constituent species-specific gene sets are simultaneously enriched in phenotype-associated genes. To test for this simultaneous enrichment, we combined enrichment p-values obtained from each gene set, then tested the significance of the combined p-value by simulating a null distribution according to a precisely specified null model.

In each species, for each orthogroup we only considered genes in the orthogroup and in the corresponding species’ gene “universe”, that is, the full complement of genes expressed above a threshold in each species. Under the null hypothesis of no orthogroup activity in response to social challenge, we modeled the number of DEGs contained in an orthogroup as a hypergeometric random variable. We tested if each orthogroup contained more DEGs from that species than expected by chance, using a one-tailed hypergeometric test. We did not separate brain region- and time point-specific DEGs within each species at this stage of analysis. This resulted in three p-values for each orthogroup, *p_bee_*, *p_mouse_*, and *P_ftsh_*, which we then aggregated using Fisher’s combination test statistic *T* = –2 ln *p_bee_* –2 ln *p_mouse_* – 2 In *p_fish_*.

For each orthogroup triplet, we calculated the p-value of the test statistic T under the reasonable assumption that the *p_bee_, p_mouse_*, and *P_fish_* were statistically independent. If they were uniformly distributed, classical theory gives that T would be 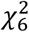-distributed under the null hypothesis that none of the three orthogroups was responsive to social challenge. However, due to the discrete nature of the hypergeometric variables from which the *p_bee_*, *p_mouse_*, and *p_fish_* were calculated, we resorted to simulations to calculate the true p-value of T. We simulated 5 million instances of the hypergeometric variables for each orthogroup in each species under the null hypothesis and calculated the p-value of each orthogroup triplet’s T using the simulated distribution.

Technically, the alternative hypothesis of this test is that there is at least one species in which the corresponding orthogroup is enriched in social challenge DEGs. This does not exactly match the conservation hypothesis, which should state that *all* orthogroups in all three species are enriched in DEGs. However, this latter hypothesis corresponds to a composite null hypothesis, which is difficult to formally test without sacrificing a great deal of statistical power. Here we instead test the simpler sharp null hypothesis where all orthogroups are inactive, it is known that the test statistic T that we have chosen is oriented toward the desired alternative hypothesis where all three orthogroups are enriched in DEGs. Thus, our tests are oriented toward the desired conservation hypothesis.

### Identifying GO terms and TF orthogroups with a conserved response to social challenge

We downloaded GO annotations for mouse and stickleback from Ensembl Biomart (Ensembl v83, Kinsella *et al*. 2011) and for bee from Ensembl Metazoa Biomart (v29, Kinsella *et al*. 2011) and considered only the 341 terms that contained at least 5 genes as HFGs. We used our orthogroup analysis method, described above, to identify terms that were significantly enriched in DEGs in multiple species. TRNs were reconstructed for each individual species individually as previously described (Bukhari *et al*. 2017; Saul *et al*. 2017; Shpigler *et al*. 2017b). In each species, for each orthogroup of TFs, we collected the gene targets of all TFs in the orthogroup into a single set. We then used our orthogroup analysis method to identify TF orthogroups whose target sets were enriched in DEGs in multiple species as HFGs.

### CNSRV

We developed a method to discover homologous gene coexpression modules across divergent species as a method of *ab initio* discovery of HFGs. This was necessitated by our multi-scale analysis strategy but may also be of independent interest. Our module inference method, called “Common NetworkS ReVealed” (CNSRV), is closest in spirit to the OrthoClust method (Yan *et al*. 2014), but uses a different score for the quality of cross-species modules. This score helps avoid a bias towards large or small modules that is commonly seen with existing methods of module discovery (Langfelder & Horvath, 2008). We performed systematic assessments to demonstrate that our methods led to less extreme module sizes (**Supplementary Figure SM2**) and also found the resulting modules to be more statistically enriched for Gene Ontology terms (**Supplementary Figure SM3**). The code for CNSRV has been deposited in GitHub (https://github.com/weiyangedward/CNSRV). For a full description of CNSRV and its evaluations, see **Supplementary Methods**. We outline its main steps below.

#### Construction of coexpression networks

For each species, we first calculated coexpression of gene pairs as the Pearson correlation of their expression values in a specific brain region at different time points after exposure (including intruder-exposed as well as control animals) and retained pairs that had correlation coefficient above 0.7 in all brain regions considered for that species.

#### Cross-species coexpression module detection

The algorithm partitions the genes in each species’ coexpression network into *K = 20* non-overlapping clusters, referred to by identifiers 1, 2, … *K*, such that cluster *i* in one species “corresponds to” clusters labeled *i* in the other species. The algorithm seeks to find partitions such that (1) clusters in each species exhibit “modularity” (Newman 2006) – high density of within-cluster coexpression edges compared to cross-cluster density of such edges, and (2) corresponding clusters in a pair of species exhibit high density of orthology edges (an orthology edge is created for any pair of genes in the same orthogroup from the two species). To meet these two goals, the CNSRV method attempts to maximize the following objective function:

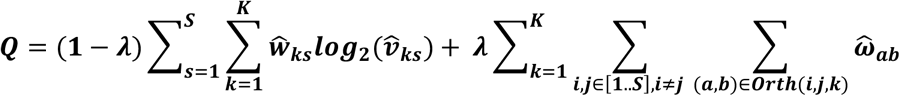

Here *S* is the number of species, *K* is the desired number of clusters. 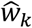 is the normalized count of coexpression edges in cluster *k* of species *s*, defined as 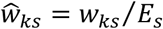 where *w_ks_* is the number of coexpression edges in cluster *k* of that species and *E_s_* is the total number of edges in that species. Similarly, 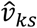 is the normalized count of coexpression edges connected to nodes in cluster *k* of species *s*, defined as 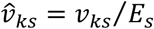, where *v_ks_* is the count of coexpression edges incident to nodes in cluster *k* in that species, (*a*, *b*) refers to any pair of orthologous genes from species *i* and *j* such that both genes are in cluster *k* of their respective species. To normalize the number of orthologous edges from many-to-many gene mappings, 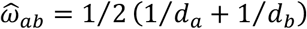 where *d_a_* is the number of orthologous edges from gene *a* in species *i* to genes in species *j*. The two terms in this formula, representing the “modularity” and “orthology” goals respectively, are weighted by factors of λ and (1 - λ) respectively. We chose a value of λ = 0.05 to provide a suitable balance between the coexpression modularity and cross-species sharing aspects of our desired gene modules (**Supplementary Figure SM1**).

The objective function is maximized with a Simulated Annealing algorithm. Initially, genes are assigned random cluster labels from 1 to *K* and the “temperature” variable is set to 10. In each proposed move, a gene is selected at random and assigned a different cluster label. The objective function is re-evaluated, “good” moves that generate a better score are accepted, whereas “bad” moves are rejected with probability that depends on the score of the proposed reassignment and the temperature variable. Specifically, the probability of accepting a proposed move that generates a new clustering with score Q_new_, assuming the current score is Q_cur_, is given by min(1, (Q_new_/Q_old_)^T^), where temperature *T* changes across iterations according to the cooling schedule *T*_*k*+1_ = α*T_k_*, where α = 0.9, and *k* is the iteration index. This results in bad moves being rejected with low probability in earlier iterations (when the “temperature” is higher), and with higher probability in later iterations. The iterative procedure stops once no good move can be found after certain amount of attempts, or a pre-determined number of iterations have been performed.

### Identifying and annotating gene coexpression modules with a conserved response to social challenge

We used 19-df chi-square tests of independence to test if the DEGs in each species/brain region/time point combination were distributed randomly across the 20 modules. A non-random distribution indicates that exposure to social challenge results in certain modules being more activated than others. To identify the active ones, we used *post-hoc* hypergeometric enrichment tests in the species/brain region/time point combinations with significant chi-square tests.

It is not clear immediately clear how to annotate these active modules in a way that also accounts for the available orthology information across the species. This is because annotations for the same module can change depending which species is considered. To address this issue, we employed a multi-species extension of our previously reported DRaWR tool (Blatti & Sinha 2016, see Supplemenatary Methods for details). DRaWR takes a heterogeneous biological network with gene and annotation nodes and ranks all annotation nodes in the network for their proximity to a set of gene nodes of interest by using random walks (**Figure 2**). We constructed a network containing “gene nodes” representing genes from all three species and “annotation nodes” that represent Gene Ontology annotations (obtained from Biomart for Ensembl v83, Kinsella *et al*. 2011) and Pfam domains (whose presence was predicted using HMMER, Finn *et al*. 2011). Edges connected genes with their properties (GO annotations and Pfam domains), and also connected homologous pairs of genes from the same or different species. For a given module, we executed the DRaWR random walk with restarts from module genes from all three species, so that the method is also able to “walk” from a gene to its ortholog(s) in other species. Separately, we also executed the random walk with restarts from module genes of each species individually, and selected annotation nodes that were ranked highest across all four restart configurations. As such, an annotation that is highly ranked by our multi-species DRaWR technique is either enriched in module genes from multiple species or enriched in orthologs of those genes (even if it is not enriched in the module genes themselves), or both. We also required that the reported annotations be significantly enriched (p-value < 0.05 using one-sided Fisher exact test) in at least one of the three species. To our knowledge, the resulting “multi-species DRaWR” algorithm is the first method capable of functional annotation of gene sets in a cross-species manner, making it ideal for identifying HFGs across distantly related species.

## RESULTS

### Brain transcriptomic response to social challenge in three diverged species shares several orthologous gene groups

We profiled gene expression by sequencing mRNA at 30 min, 60 min, or 120 min after exposure to an intruder from discrete brain regions chosen for each species: the mushroom bodies in honey bee; the amygdala, frontal cortex, and hypothalamus in mouse; and the diencephalon and telencephalon in stickleback. We considered only genes that were sufficiently expressed in these RNA-seq experiments for downstream analysis 10,701 in honey bee, 15,388 in mouse, and 17,435 in stickleback. Differentially expressed genes (DEGs) were obtained by comparison of intruder-exposed animals to control animals in matched conditions (see **Materials and Methods**), providing three sets of DEGs in honey bee, nine sets in mouse, and six sets in stickleback; these results have been reported elsewhere as individual species studies (Bukhari *et al*. 2017; Saul *et al*. 2017, Shpigler *et al*. 2017b) and are summarized in **Figure 1A**, but this is the first time that these data have been analyzed and discussed in a comparative context. These DEG sets varied in size between 36 genes (mouse amygdala, 60 min) to 1,151 (honey bee mushroom bodies, 120 min).

We were first interested in whether the same (orthologous) genes were associated with social challenge responses across these three species. However, the great evolutionary divergence of these species precludes unambiguous ortholog assignments at the gene level. We instead used orthologous groups (“orthogroups”) of genes as our fundamental unit of analysis. A resulting major analytical challenge is that most orthogroups contain different numbers of paralogs in the genomes of each individual species, and furthermore different numbers of brain regions were assessed in each species. These challenges make it difficult to ensure a fair comparison across species. Overcoming these issues requires carefully designed statistics, and existing approaches to this type of analysis, such as the one we previously employed in ref. (Rittschof *et al*. 2014), cannot be applied. To address this problem, we developed a method to identify orthogroups with the strongest evidence for activity in multiple species, where activity was measured by the proportion of DEGs, at any time point and brain region, within the orthogroup in each species (see **Materials and Methods**). Our procedure is based on another algorithm we recently developed called Orthoverlap (Shpigler *et al*. 2017a) and offers stringent control of false positives.

We obtained 4,982 orthogroups common to the three species from the OrthoDB database (Kriventseva *et al*. 2015), and our method identified six orthogroups that were responsive to a social challenge in all three species at FDR ≤ 0.10 (**Figure 3, Supplementary Table 1**). Three of the six contained at least one DEG in each of the species. Group EOG80K992 (p-value ≤ 1 × 10^−7^) – which includes the mouse genes *F5*, *Nrp2*, *Sned1*, and *Vwf* – is potentially involved in a deeply conserved immune response (Chang *et al*. 2012), but is also related to neurite outgrowth (Hey-Cunningham *et al*. 2013) and axon guidance (Klagsbrun & Eichmann, 2005). Group EOG8THX4X (p-value = 8.2 × 10^−6^), which includes the mouse gene *Npas4*, is a gene that is involved in activity-dependent development of synapses (Lin *et al*. 2008) and that regulates the balance between GABA and glutamate in neural circuits (Spiegel *et al*. 2014). This finding is consistent with our previous work (Rittschof *et al*. 2014; Saul *et al*. 2017), which also identified *Npas4* as a central gene in the shared response to social challenge based on transcriptomic analysis. Finally, group EOG8TMSCQ (p-value = 1.6 × 10^−5^) includes subunits of the heat shock protein 70 family, which is nominally associated with stressors like heat shock that require protein refolding and that often acts in concert with co-chaperones in the heat shock protein 90 family (Mayer & Bukau, 2005). Heat shock proteins from the Hsp70/Hsp90 complex have an additional documented but less discussed role, being necessary for ligand binding and subsequent signal transduction of nuclear receptors and other signaling molecules (Pratt & Toft, 2003).

**Figure 3:**
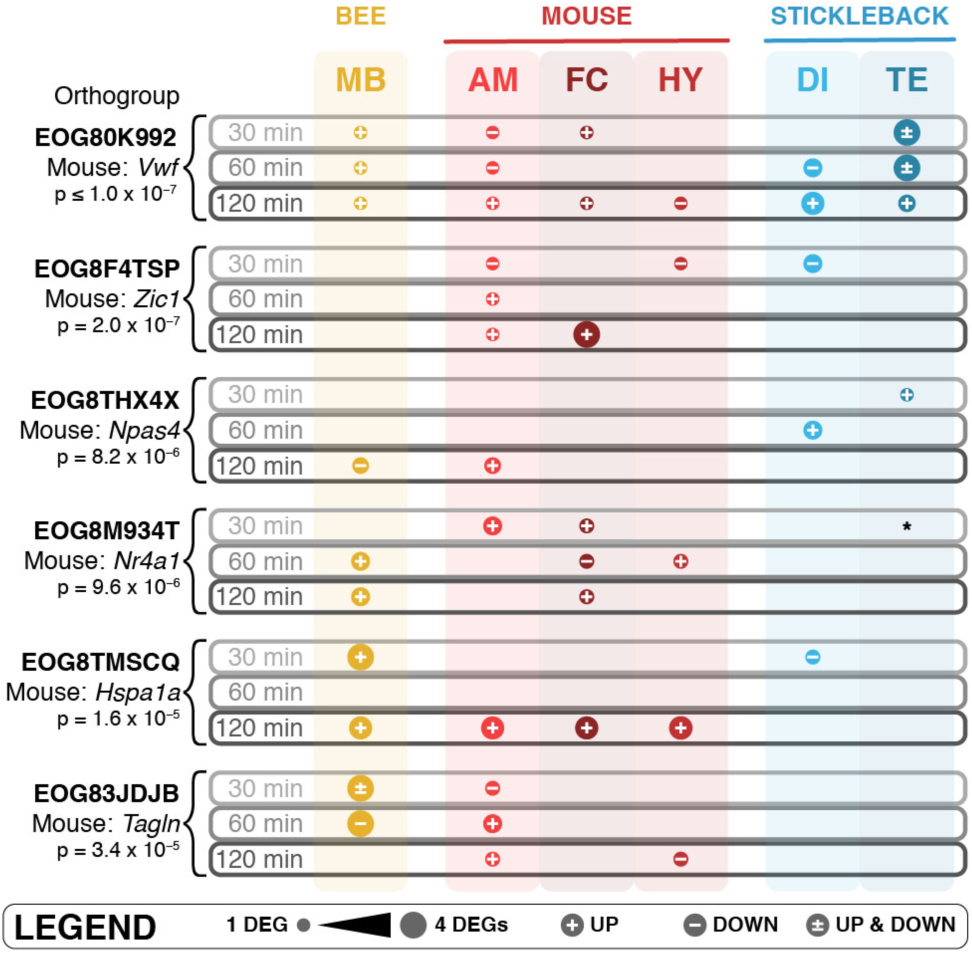
Orthogroups with significant conservation of differential expression across all three species.

The remaining three statistically significant orthogroups include DEGs in two out of the three species. We still considered these orthogroups of potential importance. For example, group EOG8M934T (p-value = 9.6 × 10^−6^), which includes the mouse gene *Nr4a1*, only contained DEGs in honey bee and mouse. However, one of the stickleback orthologs was detected at an FDR of 0.1012 (uncorrected p-value = 0.0189) in telencephalon at 30 min, only slightly higher than the 10% FDR cutoff used for that species. This group of *Nr4a* orthologs, orphan nuclear receptors with unknown ligands, thus appears to have common socially regulated activity. These receptors, which are known to regulate glucose metabolism and homeostasis (Close *et al*. 2013), have documented roles in memory and in object recognition (McNulty *et al*. 2012) and have been documented as related to social aggression in vertebrates previously (Malki *et al*. 2016). Additionally, group EOG8F4TSP (p-value = 2 × 10^−7^), which contains “zinc finger of the cerebellum” (Zic) proteins, contains at least one DEG in both mouse and stickleback, but not in honey bee. This group of C2H2 zinc finger proteins is known for their evolutionarily conserved roles in neural development (Aruga, 2004; Fujimi *et al*. 2006).

Several of these findings rose to significance only because we collected RNA-seq data in three species. For example, *Npas4* and *Nr4a1*, transcription factors involved in neural function and/or development, had not been identified as central molecules in the response to social challenge in each individual species (but see Rittschof *et al*. 2014; Shpigler *et al*. 2017b), but our comparative analysis showed that these genes were consistently involved in the brain’s response to challenge in all three of our species. The multiple brain region/time point resolution of these RNA-seq data also allowed us to identify shared genes that are transiently expressed, and/or expressed in a brain-region specific manner. For example, the heat shock orthogroup, which contains the chaperone gene *Hspa1a*, a potential cofactor with nuclear receptors like *Nr4a1*, was only active at 120 min in the mouse and in the diencephalon in the stickleback and would have been missed if not for the time course design.

### Social challenge triggers shared hormone-dependent neuronal signaling

Shared mechanisms of the response to social challenge may emerge at higher levels of organization than that of individual genes. We asked if the same cellular processes (e.g., Gene Ontology terms) are transcriptionally active in response to social challenge, even if specific genes exhibiting differential expression are not strictly orthologs of each other. This allows us to be more sensitive to cellular mechanisms that may have evolved by convergence through repeated coopting the same biological pathways.

We considered gene sets defined by 341 GO terms that contained at least 5 genes in each of the species studied here. Using the method that we developed for our analysis of shared gene orthogroups above, we identified those GO terms that had the strongest evidence for enrichment of DEGs in each of the three species. We identified 66 GO terms at FDR ≤ 0.10 and 37 GO terms at a more stringent threshold of FWER ≤ 0.10 (**Table 1**). These terms centered on five major categories: hormone activity, transmembrane transport, G-protein coupled signal transduction, synaptic activity, and extracellular matrix components. This analysis thus identified a set of processes that correspond to the same general functions, though they have a slightly different complement of genes between distantly related taxa. Specifically, these results suggest that hormone receptors, acting as nuclear receptors, signaling molecules and transcription factors, are essential in the coordination of the large-scale social challenge induced transcriptional responses that potentially cause remodeling of axons and dendrites, which lead to differences in synapse-related proteins, extracellular matrix proteins, transmembrane transporters, and the modulation of GPCRs for neural signaling.

**Table 1:** Multiple cross-species-mapped GO terms show conserved activity in response to social challenge (includes Biological Processes, Cellular Components, and Molecular Functions).

| Biological Process GO ID – Term | P Sim. | Ratio (Hits / Total Genes in GO Term) |  |  |
| --- | --- | --- | --- | --- |
|  |  | Honey Bee | Mouse | Stickleback |
| GO:0007186 – G-protein coupled receptor signaling pathway | $< 1 \times 10^{-7}$ | 30/139 | 52/344 | 58/399 |
| GO:0007218 – neuropeptide signaling pathway | $< 1 \times 10^{-7}$ | 5/17 | 20/70 | 2/6 |
| GO:0007601 – visual perception | $< 1 \times 10^{-7}$ | 1/6 | 7/71 | 12/16 |
| GO:0055085 – transmembrane transport | $< 1 \times 10^{-7}$ | 36/246 | 42/338 | 67/425 |
| GO:0007155 – cell adhesion | $2.0 \times 10^{-7}$ | 8/54 | 40/308 | 20/120 |
| GO:0006836 – neurotransmitter transport | $4.0 \times 10^{-7}$ | 4/16 | 11/28 | 3/28 |
| GO:0043401 – steroid hormone mediated signaling pathway | $5.4 \times 10^{-7}$ | 11/21 | 9/52 | 5/61 |
| GO:0007169 – transmembrane receptor protein tyrosine kinase signaling pathway | $5.8 \times 10^{-6}$ | 1/7 | 14/81 | 11/45 |
| GO:0006811 – ion transport | $6.6 \times 10^{-6}$ | 14/108 | 17/176 | 39/190 |
| GO:0006366 – transcription from RNA polymerase II promoter | $7.2 \times 10^{-6}$ | 2/22 | 41/314 | 0/6 |
| GO:0007165 – signal transduction | $1.8 \times 10^{-4}$ | 50/285 | 60/633 | 50/506 |
| GO:0007166 – cell surface receptor signaling pathway | $2.9 \times 10^{-4}$ | 3/16 | 17/132 | 11/58 |

| Cellular Component GO ID – Term | P Sim. | Ratio (Hits / Total Genes in GO Term) |  |  |
| --- | --- | --- | --- | --- |
|  |  | Honey Bee | Mouse | Stickleback |
| GO:0005576 – extracellular region | $< 1 \times 10^{-7}$ | 27/113 | 85/481 | 31/176 |
| GO:0005578 – proteinaceous extracellular matrix | $< 1 \times 10^{-7}$ | 2/20 | 40/193 | 3/14 |
| GO:0005615 – extracellular space | $< 1 \times 10^{-7}$ | 6/17 | 107/712 | 5/18 |
| GO:0005886 – plasma membrane | $< 1 \times 10^{-7}$ | 8/58 | 203/2146 | 23/109 |
| GO:0005887 – integral component of plasma membrane | $< 1 \times 10^{-7}$ | 3/7 | 83/562 | 1/9 |
| GO:0016021 – integral component of membrane | $< 1 \times 10^{-7}$ | 119/785 | 245/3266 | 153/1212 |
| GO:0016459 – myosin complex | $< 1 \times 10^{-7}$ | 8/23 | 2/39 | 21/53 |
| GO:0031012 – extracellular matrix | $2.0 \times 10^{-7}$ | 2/11 | 28/158 | 6/40 |
| GO:0016020 – membrane | 4.8 x 10<sup>-6</sup> | 123/813 | 147/2521 | 174/1229 |
| GO:0045202 – synapse | 6.7 x 10<sup>-5</sup> | 1/25 | 31/236 | 1/6 |

| Molecular Function GO ID – Term | P Sim. | Ratio (Hits / Total Genes in GO Term) |  |  |
| --- | --- | --- | --- | --- |
|  |  | Honey Bee | Mouse | Stickleback |
| GO:0003774 – motor activity | < 1 x 10<sup>-7</sup> | 8/23 | 2/51 | 21/53 |
| GO:0004930 – G-protein coupled receptor activity | < 1 x 10<sup>-7</sup> | 26/124 | 37/247 | 51/360 |
| GO:0005179 – hormone activity | < 1 x 10<sup>-7</sup> | 3/8 | 12/41 | 10/40 |
| GO:0005509 – calcium ion binding | < 1 x 10<sup>-7</sup> | 29/169 | 57/488 | 90/535 |
| GO:0043565 – sequence-specific DNA binding | 2.0 x 10<sup>-7</sup> | 32/133 | 44/385 | 48/365 |
| GO:0005515 – protein binding | 1.2 x 10<sup>-6</sup> | 279/1635 | 534/8766 | 417/3710 |
| GO:0003707 – steroid hormone receptor activity | 1.4 x 10<sup>-6</sup> | 10/18 | 9/45 | 5/62 |
| GO:0005216 – ion channel activity | 2.4 x 10<sup>-6</sup> | 5/65 | 11/109 | 30/119 |
| GO:0005198 – structural molecule activity | 4.8 x 10<sup>-6</sup> | 2/24 | 12/86 | 20/79 |
| GO:0005201 – extracellular matrix structural constituent | 5.6 x 10<sup>-6</sup> | 4/7 | 6/26 | 7/25 |
| GO:0020037 – heme binding | 3.3 x 10<sup>-5</sup> | 10/63 | 15/77 | 11/72 |
| GO:0004714 – transmembrane receptor protein tyrosine kinase activity | 3.4 x 10<sup>-5</sup> | 1/6 | 8/36 | 8/28 |
| GO:0005215 – transporter activity | 4.6 x 10<sup>-5</sup> | 13/95 | 19/110 | 16/122 |
| GO:0005506 – iron ion binding | 2.6 x 10<sup>-4</sup> | 8/60 | 16/95 | 11/82 |
| GO:0003700 – transcription factor activity, sequence-specific DNA binding | 2.7 x 10<sup>-4</sup> | 40/168 | 51/637 | 38/346 |

### Shared gene coexpression modules responding to social challenge

In addition to defining sets of genes using GO terms, we sought to identify coordinately expressed sets of genes, often called gene modules, *ab initio*, without the need for prior knowledge. Coexpressed gene modules have become a mainstay of systems-level analysis of transcriptional programs (Langfelder & Horvath 2008). Studies in evolutionary developmental biology have noted that modules underlying development are deeply conserved and are an important facet of the genetic “toolkit” concept (Peter & Davidson 2011; Rittschof & Robinson 2016; Toth & Robinson 2007). Coexpressed modules are also conserved in other biological contexts across evolutionary spans as great as humans, flies, worms, and yeast (Stuart *et al*. 2003), but typically have not been explicitly examined as an HFG representing shared responses to social stimuli.

Using the new developed CNSRV method for cross-species analysis, we sought to discover broadly shared gene modules (see schematic representation of coexpressed shared modules in **Figure 4A**) from our multi-species brain transcriptomic data, then query if any of these are regulated by social challenge commonly across the three species. Our experimental design allowed us to characterize modules with coordinated anatomic and temporal expression profiles in bee, mouse, and stickleback. We then combined these results with our DEGs to identify modules that were highly responsive to social challenge (see **Materials and Methods**). These represent shared core regulatory programs, where conservation is identified at the level of coexpressed modules rather than individual genes.

**Figure 4:**
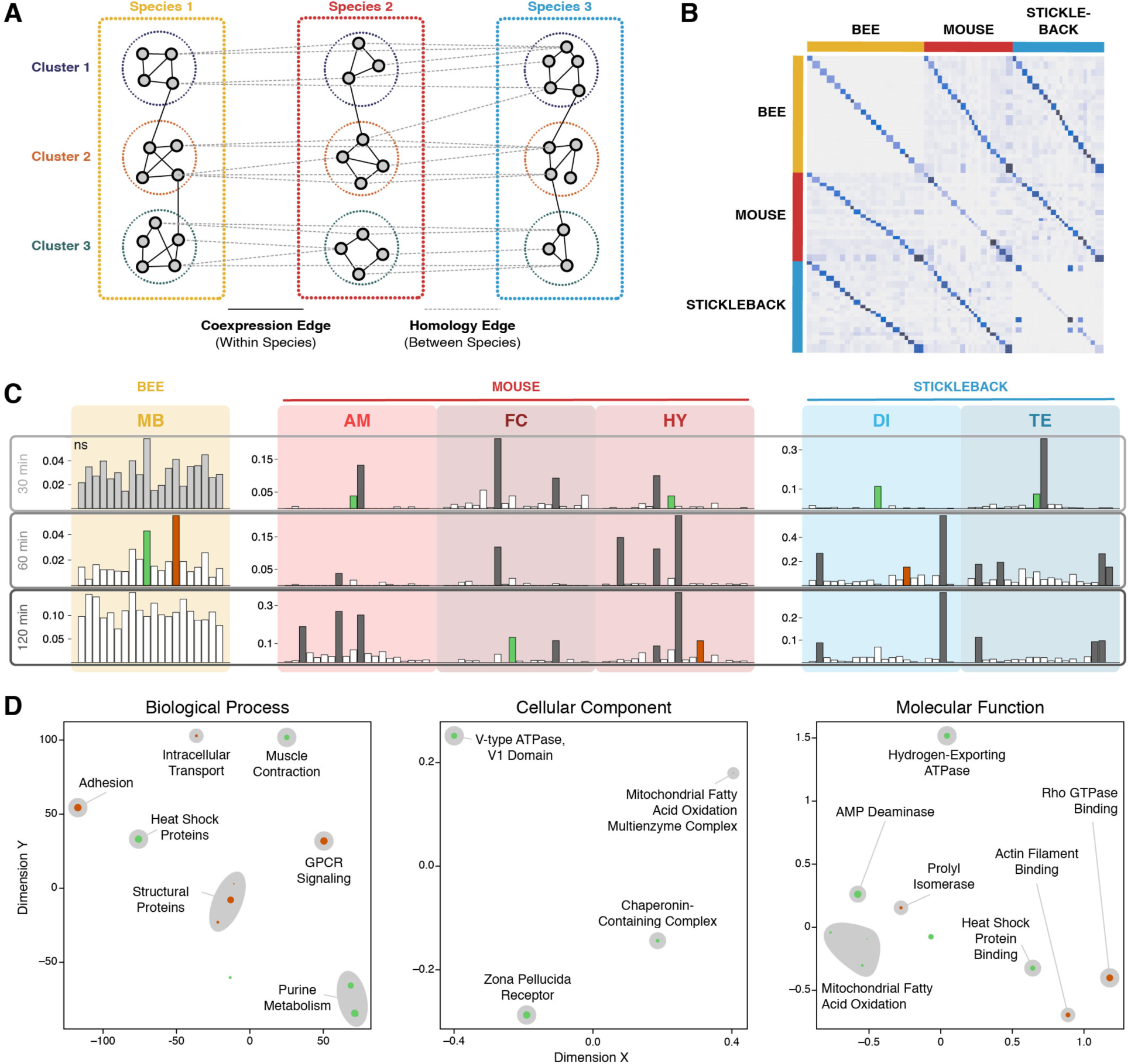
Cross-species coexpression module algorithm (**A**) conceptual schematic and (**B-D**) results. **A**) Schematic of CNSRV, the cross-species clustering algorithm used to find conserved gene modules, which uses evidence derived from both coexpression and conservation to find gene modules enriched in conservation. **B**) Clustering results from CNSRV show that conserved modules, shown by the ancillary diagonals off the main diagonal, cluster better between species than do unmatched modules. **C**) Enrichment results for DEGs for CNSRV modules within each species reveal significant differences among clusters in all but honey bee mushroom body at 30 min (light gray). Multiple CNSRV clusters were enriched in individual species (dark gray), but two modules – 10 and 14, shown in green and dark orange respectively – show simultaneous enrichment for differential expression across all 3 species. **D**) Multidimensional scaling on semantic distances for GO terms enriched in the multi-species DRAWR results show clusters of GO terms commonly related to clusters 10 and 14 across all 3 species. Larger points associated with each GO term correspond to stronger p-values. Gray clouds correspond to a high-level biological description of the GO terms within each cluster annotated by the authors.

With CNSRV, we identified 20 homologous modules (**Figure 4B**), each ranging between 140 and 523 genes in size (see **Materials and Methods** and **Supplementary Table 2**). These modules show both dense coexpression within modules in individual species (Figure 2B, central diagonal) and elevated frequency of orthology relationships between corresponding modules (Figure 2B, ancillary diagonals). Next, for each combination of brain region and time point in each species, we tested if DEGs were differentially distributed among the modules (see **Materials and Methods**) and found this to be the case (FDR ≤ 0.10) for all but one of the 18 species/brain region/time point combination (**Figure 4C**). Within these 17 significant combinations of region, time, and species, we then conducted *post-hoc* tests at FWER ≤ 0.10 to identify significantly enriched modules.

**Figure 2:**
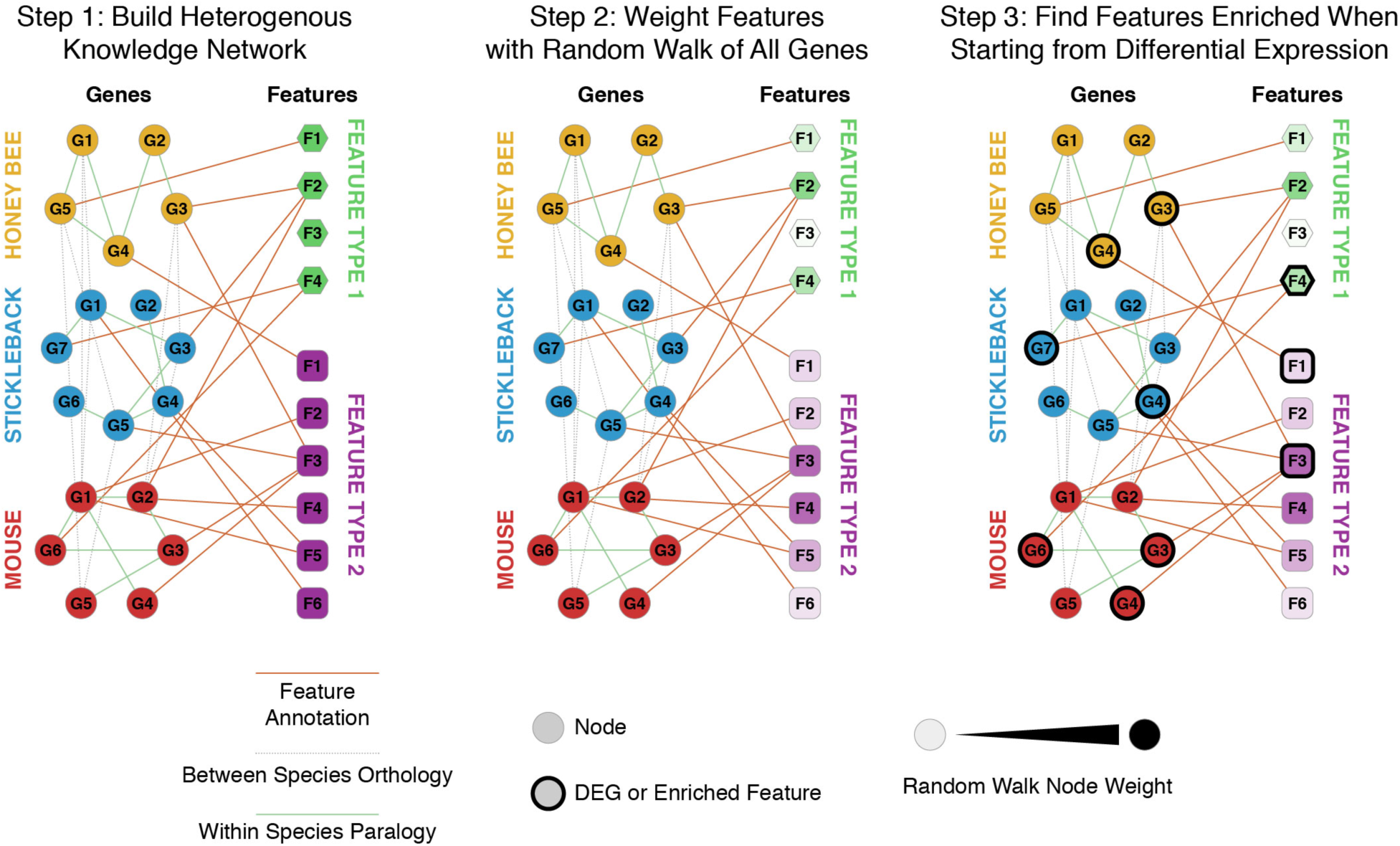
Schematic of heterogeneous knowledge network enrichment using cross-species DRaWR. In brief, the algorithm operates in 3 steps: **1**) Construct a heterogeneous network consisting of edges connecting gene nodes with one another within species (paralogy relationships in this dataset), with one another between species (cross-species orthology information in this dataset), and with annotations features (Gene Ontology terms and Pfam domains in this dataset). **2**) Using a random walk with restarts starting on the whole gene set, identify features that are frequently traversed and weight them appropriately. **3**) Using a random walk with restarts on the differential expression gene set, identify features that are more frequently traversed than in all genes.

This analysis revealed that two gene coexpression modules, numbered 10 and 14, have shared social challenge-specific activity across all three species (**Figure 4C**). Specifically, module 10 is significantly associated with DEGs in honey bee mushroom body (60 min), mouse frontal cortex (120 min) and hypothalamus (30 min), as well as stickleback diencephalon (30 min) and telencephalon (30 min). Similarly, module 14 is enriched for DEGs in honey bee mushroom body (60 min), mouse hypothalamus (120 min), and stickleback diencephalon (60 min). We note that time points where the orthologous modules were observed often did not match between the species, which may have resulted from differences in the timing of the neurobiological responses of each species, underscoring the importance of multiple time points in the study design.

While it was instructive to observe shared modules apparently regulated by social challenge, it was not as clear what biological functions these modules might be involved with. Functional annotation of these modules is difficult because a module in our context is not a single list of genes but a set of three different species-specific gene lists with strong mutual orthology, and standard gene set enrichment tests do not take into account the multi-species nature of the modules or their orthology relationships. To solve this problem, we adapted our previously developed tool DRaWR (Blatti & Sinha, 2016), which considers a network whose nodes are genes and annotations (e.g., Gene Ontology terms) and edges connect a gene to each of its annotations. It annotates a gene set by performing a random walk starting from nodes in the gene set and recording the annotation nodes that are visited most frequently. We extended this approach here to annotate the orthologous CNSRV modules by constructing a network using module genes, GO annotations, and orthology edges from all three of our species (see **Materials and Methods**).

**Figure 4D** shows the top functional annotations for modules 10 and 14, as revealed by a high DRaWR percentile score in every species, and with additional support from standard enrichment tests (hypergeometric test nominal p-value ≤ 0.05) in at least two of the three species. Module 14 comprises genes involved in cell-matrix adhesion, a process involved in neural development and plasticity (Murase & Schuman 1999); Rho GTPase binding, a process implicated in several aspects of neuronal development as well as neurological diseases (Govek *et al*. 2005); and actin binding, a process associated with function and plasticity of dendritic spines and synapses (Lin & Webb, 2009). Module 10 includes genes annotated for AMP deaminase activity and IMP biosynthesis, processes associated with purine balance in the brain. Purine balance and purinergic reception play well-known roles in neuronal repair and protection, acting as a bridge between neural signaling and the neural immune system in mammals (Skaper *et al*. 2010; Thauerer *et al*. 2012). Further, the enrichment of enoyl-CoA hydratase activity found in Module 10, as a step of fatty acid metabolism found in the Cellular Component and Molecular Function results, potentially bridges neural signaling and the metabolic processes previously observed in response to social challenge both across species and within individual species (Chandrasekaran *et al*. 2015; Rittschof *et al*. 2014). These coexpression modules bolster evidence from the shared DEGs and from the GO results in support of a shared transcriptomic response that includes structural proteins, heat shock proteins, and GPCR signaling proteins.

### Common transcription factor regulatory activities underlie social challenge

The previous sections provide insight into the common biological processes and gene modules underlying the response to social challenge. We next sought to identify transcription factors (TFs) that act as master regulators of those processes and modules, using state-of-the-art tools for reconstruction of transcriptional regulatory networks in each species. In particular, we queried if the same TFs (or their paralogs) regulate the brain transcriptomic response to social challenge across species. TF-gene relationships are among the best studied and most widely accepted conception of gene networks, and they have been explored in the context of genetic toolkit studies in evo-devo (Rittschof & Robinson, 2016). The gene orthogroup analysis reported above (**Figure 3**) identified multiple TF orthogroups containing social challenge DEGs; however, there may be TFs which do not detectably change in transcript expression upon social challenge but may for example be activated by post-transcriptional modifications. Regulatory targets for these TFs may nevertheless be socially regulated and the TFs reasonably speculated to have a role in the transcriptional response to social challenge. We reasoned that even if the TFs are not differentially expressed on social challenge, their regulatory targets may still be identifiable from a covariance between TF and gene expression across the many brain regions and time points assayed here, allowing us to test enrichment between each TF’s targets and social challenge DEGs.

To explore this idea, we constructed transcriptional regulatory networks (TRNs) for each species using the previously developed tool ASTRIX (Chandrasekaran *et al*. 2011), which uses the ARACNE algorithm (Margolin *et al*. 2006) to identify putative TFs for a gene, then employs Least Angle Regression (Efron *et al*. 2004) to identify those TFs that best predict expression levels of that gene target in multiple experiments. In this case, each TRN had been reconstructed from different brain regions and time points within each individual species previously (Bukhari *et al*. 2017; Saul *et al*. 2017; Shpigler *et al*. 2017b). We used these previously reconstructed TRNs to identify TF orthogroups whose gene targets were enriched in DEGs in all three species, using the same method as described above for identifying the HFGs of shared gene orthogroups and GO terms. We considered only orthogroups that contained at least one TF with at least one gene target in each species. This analysis detects TFs important to social challenge even if the TFs themselves are not significantly differentially expressed.

We detected six TF orthogroups (FDR ≤ 0.10) that are likely to be conserved regulators of the transcriptomic response to social challenge (**Table 2**). For instance, the orthogroup EOG8KWM99 comprises the mouse TF genes *Pbx1* and *Pbx3*, for which the ASTRIX-derived TRN included 2 target genes in mouse, both of which are social challenge DEGs, 45 targets (including 3 DEGs) in stickleback and 25 targets (including 9 DEGs) in honey bee. Further, one orthogroup that includes the mouse neural development transcription factor genes *Rax* and *Pax6* may be involved in the conserved regulation of the formation of new neurons from a neural stem cell lineage (Davis *et al*. 2003; Pak *et al*. 2014). *Rax* was identified as a transcriptional regulator in our earlier work (Rittschof *et al*. 2014). One particularly interesting TF, the orphaned nuclear receptor mouse gene *Nr2e1*, has been implicated in our previous cross-species work (Rittschof *et al*. 2014), in aggression in mice (Abrahams *et al*. 2005), and in aggression in flies (Davis *et al*. 2014). These results identify specific transcriptional regulators that appear to be important central regulators of the processes described in the above sections and therefore constitute potential key shared master regulators of the transcriptional response to social challenge.

**Table 2:** Conserved transcription factor expressed in the brain implicated as regulators of response to social challenge

| Orthogroup | Ratio (Hits / Total Targets) |  |  | p | Mouse Gene Names |
| --- | --- | --- | --- | --- | --- |
|  | Honey Bee | Mouse | Stickleback |  |  |
| EOG8JT1HM | 1/3 | 12/84 | 30/128 | 0 | *Sp3, Klf16, Klf13, Klf10, Sp4, Klf4, Klf9, Sp1, Sp7, Klf5, Klf12, Klf7* |
| EOG873R3N | 0/1 | 8/42 | 11/34 | 0.000004 | *Barx2, Nkx2-1, Hmx2, Hhex* |
| EOG8KWM99 | 9/25 | 2/2 | 3/45 | 0.0003072 | *Pbx3, Pbx1* |
| EOG86DNH2 | 5/9 | 0/1 | 1/1 | 0.0012816 | *Nr2e1* |
| EOG8JWWWP | 4/5 | 1/6 | 0/6 | 0.0055148 | *Gsx1* |
| EOG81RRB5 | 5/105 | 1/2 | 9/42 | 0.0171374 | *Arx, Pax6, Rax* |

## DISCUSSION

The evolution of gene regulatory programs is a subject of long-standing interest (Halfon & Michelson 2002) and has been studied by cross-species comparisons of cis-regulatory sequences (Sinha *et al*. 2004), TF-DNA binding (ENCODE Project Consortium 2012), as well as gene expression measurements in matched tissues and organs (Breschi *et al*. 2017; Gerstein *et al*. 2014; Lin *et al*. 2014). An important feature of our study was its explicit coupling of gene and gene network comparisons with objectively defined and analogous phenotypic states measured experimentally to identify shared mechanisms that may constitute a behavioral toolkit. We note that such an evolutionary toolkit, consisting of a non-unitary system of genes, does not require framing in the dichotomy of common descent versus convergence. It is instead possible that shared functions derive from redeployment and elaboration of ancestral gene modules into new contexts as has been observed in insect gene regulatory network evolution (Kazemian *et al*. 2014, Suryamohan *et al*. 2016).

Our approach utilizes an array of novel tools with a common theme of studying different tiers of organization for evidence of a shared genetic program: each test assays if groups of related genes – orthogroups, functional systems, coexpressed modules, or transcription factor regulons – have a non-random association with socially responsive genes expressed in the brain in multiple species. Our methodology for cross-species associations between HFGs and phenotypes can also be used in other contexts to identify similar broadly shared molecular systems in association with other phenotypes of interest. Moreover, the scope of such transcriptome-wide comparisons distinguishes this work from more directed studies of regulatory evolution where expression and cis-regulatory divergence of individual genes was linked to morphological differences between species (Wray 2007). Our goal is similar to the work of (Malki *et al*. 2016), who compared aggression-related DEGs in brains of mouse and zebrafish, but our study pursues the goal through an experiment design wherein data were collected from multiple species in a parallel manner, addressing several key technical and statistical challenges in the process.

One technical challenge observed in these data is the difficulty in matching gene expression sets across such long evolutionary distances. We note that time points where the orthologous modules were observed often did not match between the species, which may have resulted from differences in metabolic rates between the species. This observation underscores the importance of time series in the study design. Furthermore, it demonstrates that experimental designs in future studies must proceed carefully to identify matching expression sets across species. We were able to go beyond identification of differentially expressed genes and rigorously analyze coexpression relationships only because our experimental design included multiple brain regions and time points. Thus, the design gave us access to the higher order systems mechanisms mentioned above, significantly elaborating upon our earlier work (Rittschof *et al*. 2014).

Integrating results from these multiple analytical levels, we propose a system of genes acting commonly in the adult brains of these diverged species to transduce social challenge stimuli into transcriptional and epigenomic responses. This is graphically summarized in **Figure 5**. It involves the integration of nuclear receptor signaling to drive the transcriptional regulatory events that result in changes in neural signaling observed after a social challenge. We speculate that because nuclear receptors are both liganded receptors and transcription factors, they act as key drivers of the large-scale transcriptional changes seen across all of these species. We further speculate that these transcriptional changes occur in concert with transcription factors commonly associated with neural development to drive neural signaling modulation, which likely take place through alterations in dendritic architecture, axon architecture, signaling molecules like GPCRs and ion channels, mitochondrial metabolism, or all of these processes simultaneously.

**Figure 5:**
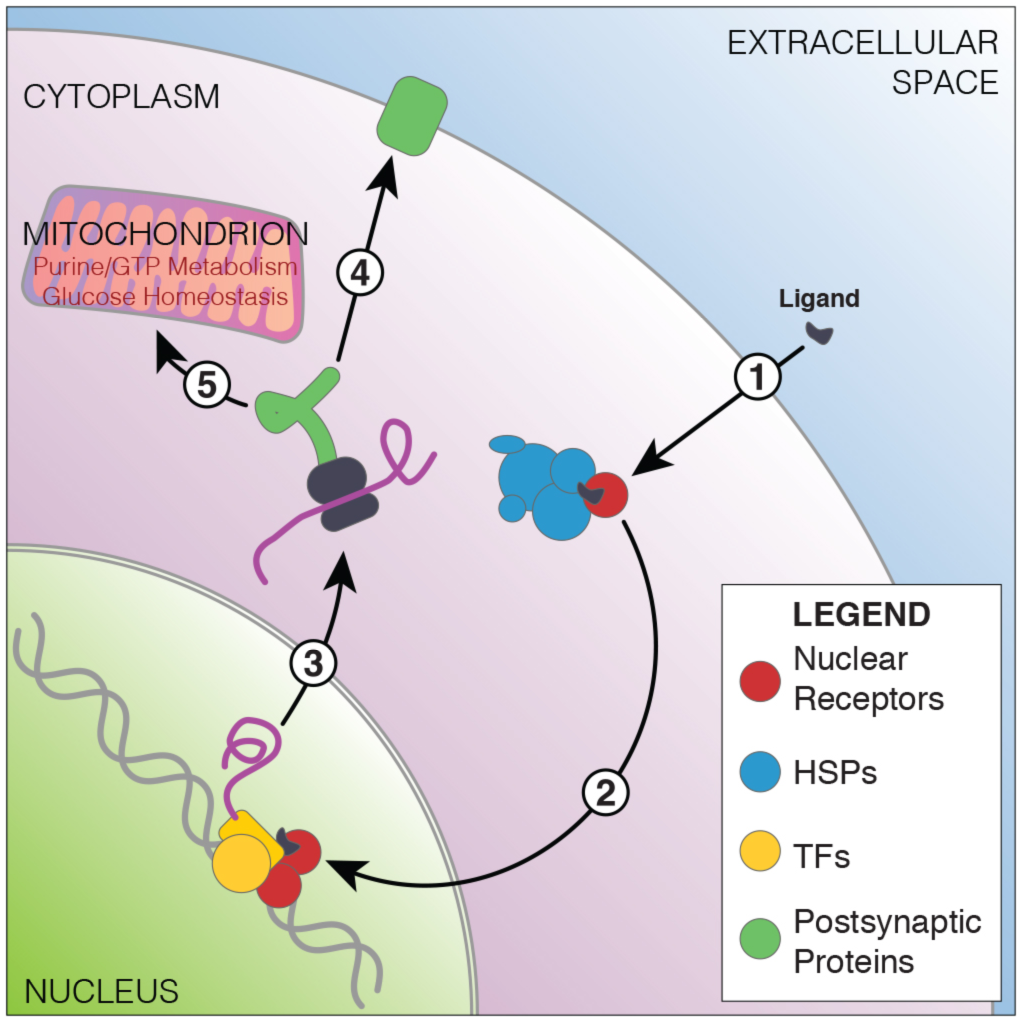
Schematic representation of genes and gene sets found enriched in the brain’s response to social challenge across honey bees, stickleback fish and mice. Hypothesized pathway includes **1**) nuclear receptor signaling interacting with heat shock/chaperones. **2**) These nuclear receptors translocate across the membrane, interacting with well-known neurally active transcription factors to cause **3**) alterations in transcription. These induce changes in **4**) postsynaptic proteins and **5**) mitochondrial function.

In this pathway, we call specific attention to the signaling molecules, transcription factors, and nuclear receptors that can act as both. Specifically, the various homologs of *Npas4*, *Nr2e1*, and *Nr4a1* are transcription factor genes well-known in neural response to stimuli (Abrahams *et al*. 2005, Kim *et al*. 2010; Maxwell & Muscat 2006). We speculate that the ancestral versions of these genes, which were likely present in the most recent common ancestor of all living bilaterians, were potentially already active in the response to social challenge stimuli that was exhibited by their contemporaries around the time of the Cambrian explosion (Carbone & Narbonne 2014). Translating these gene expression patterns into knowledge about how the cellular systems inside the brain change in response to social challenge is an important next step. Such research will require careful work across species to identify important points of similarity as well as how these systems diverge.

Though we discussed the role and neurobiological relevance of some of the above-mentioned systems in detail in our previous work – we described hormone receptors in sticklebacks (Bukhari *et al*. 2017), developmental transcription factors in mice (Saul *et al*. 2017), dendritic architecture in honey bees (Shpigler *et al*. 2017b), and GPCRs in all three species (Bukhari *et al*. 2017; Saul *et al*. 2017; Shpigler *et al*. 2017b) – the present work unifies these systems in their role in social responsiveness into a whole. The genes and systems-level mechanisms we proposed here as drivers of the response to social challenge constitute real, testable connections about a possibly conserved genetic program for the response to a social challenge, something that was lacking before this analysis. However, we note that these mechanisms may not be specific to social contexts, but may instead coordinate information from multiple contexts, and thus, the specificity of these gene sets for social challenge response also needs rigorous testing.

## ACKNOWLEDGEMENTS

We would like to thank Yoshitsugu Oono and Jian Ma for their critical comments during the compilation of this manuscript. This research was primarily funded by grant #SFLife 291812 from the Simons Foundation. In addition, this research was supported in part by grant 1U54GM114838 awarded by NIGMS through funds provided by the trans-NIH Big Data to Knowledge (BD2K) initiative and by NSF grant DMS-1613005 to SDZ.

## SUPPLEMENTARY MATERIALS

**Supplementary Methods** (Supplementary_Methods.pdf): Detailed materials and methods for the CNSRV cross-species coexpression module discovery algorithm (includes **Supplementary Figures SM1-3**).

**Supplementary Table 1** (Supplementary_Table_1.xls): Orthogroups displaying significant cross-species enrichment in DEG lists.

**Supplementary Table 2** (Supplementary_Table_2.xls): CNSRV-derived cross-species coexpression modules.

